# Breasi-CRISPR: an efficient genome editing method to interrogate protein localization and protein-protein interactions in the embryonic mouse cortex

**DOI:** 10.1101/2022.02.02.478837

**Authors:** Brandon L. Meyerink, KC Pratiksha, Neeraj K. Tiwari, Claire M. Kittock, Abigail Klein, Claire Evans, Louis-Jan Pilaz

## Abstract

In developing tissues, knowing the localization and interactors of proteins of interest is key to understanding their function. This can be challenging when the researched protein lacks reliable antibodies. Here, we combine Easi-CRISPR with *in utero* electroporation to tag endogenous proteins within embryonic mouse brains. This method is called Breasi-CRISPR (Brain Easi-CRISPR), and enables knock-in of both short and long epitope tag sequences in genes of interest with high efficiency. Using Breasi-CRISPR, we were able to visualize epitope tagged proteins known to have either high or low expression levels, such as ACTB, LMNB1, EMD, FMRP, NOTCH1, and RPL22. Detection was possible by immunohistochemistry as soon as one day after electroporation at embryonic day 13 (E13). Two and five days after electroporation, we observed efficient gene editing in up to 50% of electroporated cells. Moreover, tagged proteins could be detected by immunoblotting in lysates from individual cortices two days after electroporation. Next, we demonstrated that Breasi-CRISPR enables the tagging of proteins with fluorophores in an efficient manner, allowing the visualization of endogenous proteins via live-imaging in organotypic brain slices two days after electroporation. Finally, we used Breasi-CRISPR to perform co-IP mass-spectrometry analyses of tagged autism-related protein FMRP to discover its interactome in the embryonic cortex. Together, these data show Breasi-CRISPR is a powerful tool with diverse applications that will propel the understanding of protein function in neurodevelopment.

## Introduction

Brain development involves a myriad of mechanisms and pathways, which heavily rely on proteins operating in defined subcellular compartments, via interactions with other molecules. The study of these mechanisms classically involves analyses such as immunohistochemistry (IHC) to characterize protein localization, and immunoprecipitation approaches to probe for protein interactors. However, studying endogenous proteins in the developing cortex remains challenging. This is especially the case when a protein of interest lacks specific antibodies to perform the aforementioned analyses. Overexpression of tagged proteins in subsets of cells is used to circumvent this issue; however, protein overexpression can have drastic consequences on the cells of interest, preventing the experimenter from studying a protein in a physiological setting (Kintaka et al., 2016, Kafri et al., 2016). Moreover, having too much of a particular protein can lead to nonspecific binding to proteins with which it normally does not interact.

CRISPR technology has given researchers the opportunity to manipulate endogenous proteins to interrogate their function. In the developing mouse cortex, *in utero* electroporation (IUE) can be used to deliver reagents necessary for CRISPR-based epitope tagging (Tsunekawa et al., 2016, Suzuki et al., Mikuni et al., 2016, Uemura et al., 2016, Fang et al., 2021). Some approaches rely on the homology-directed repair (HDR) machinery operating in early progenitor cells (embryonic day 12-13 [E12-E13]) (Mikuni et al., 2016, Tsunekawa et al., 2016, Uemura et al., 2016), while others leverage the non-homologous end joining (NHEJ) machinery in late progenitor cells and post-mitotic neurons (E15 and beyond, (Suzuki et al., 2016, Fang et al., 2021)). For all these paradigms, CRISPR reagents consist of bi-cistronic expression vectors encoding a single guide RNA (sgRNA) and CRISPR-CAS9 protein, together with a separate DNA fragment to direct HDR or NHEJ to introduce a sequence of interest. Here, we are coupling IUE in the developing mouse cortex with the Easi-CRISPR approach (Efficient additions with ssDNA inserts-CRISPR, (Miura et al., 2018, Quadros et al., 2017). Easi-CRISPR has mainly been used to generate transgenic mice by electroporating single cell embryos with pre-formed RNP complexes composed of recombinant CAS9 and synthetic guide RNAs together with a chemically enhanced single-stranded oligo donor (Fig. 1A). We have moved this approach to the developing brain, electroporating neuronal precursors with these RNP complexes to selectively edit neural clonal lineages. We call this approach Breasi-CRISPR (Brain Easi-CRISPR). We report that Breasi-CRISPR is an efficient and rapid method to introduce epitope-tag sequences in cells of the developing cortex, enabling the visualization of endogenous protein in individual cells as soon as one day after electroporation in up to 50% of the electroporated cells, and in 30% of all the cells of the electroporated region. However, the more significant technical advance provided by this approach is that the high efficiency of recombination enables the detection of tagged proteins by immunoblot analyses in single cortices in as little as two days after electroporation and by IHC in as little as 24 hours. To further demonstrate the power of this approach, we tagged endogenous proteins with EGFP to visualize them by live-imaging in brains slices two days after IUE. Finally, we performed co-IP mass-spectrometry experiments with an endogenously epitope-tagged protein. These experiments led to the detection of protein-protein interactions consistent with the previously known role of the tagged protein. Thus, this technique can be used to interrogate function of endogenous proteins during early brain development.

**Figure 1.**
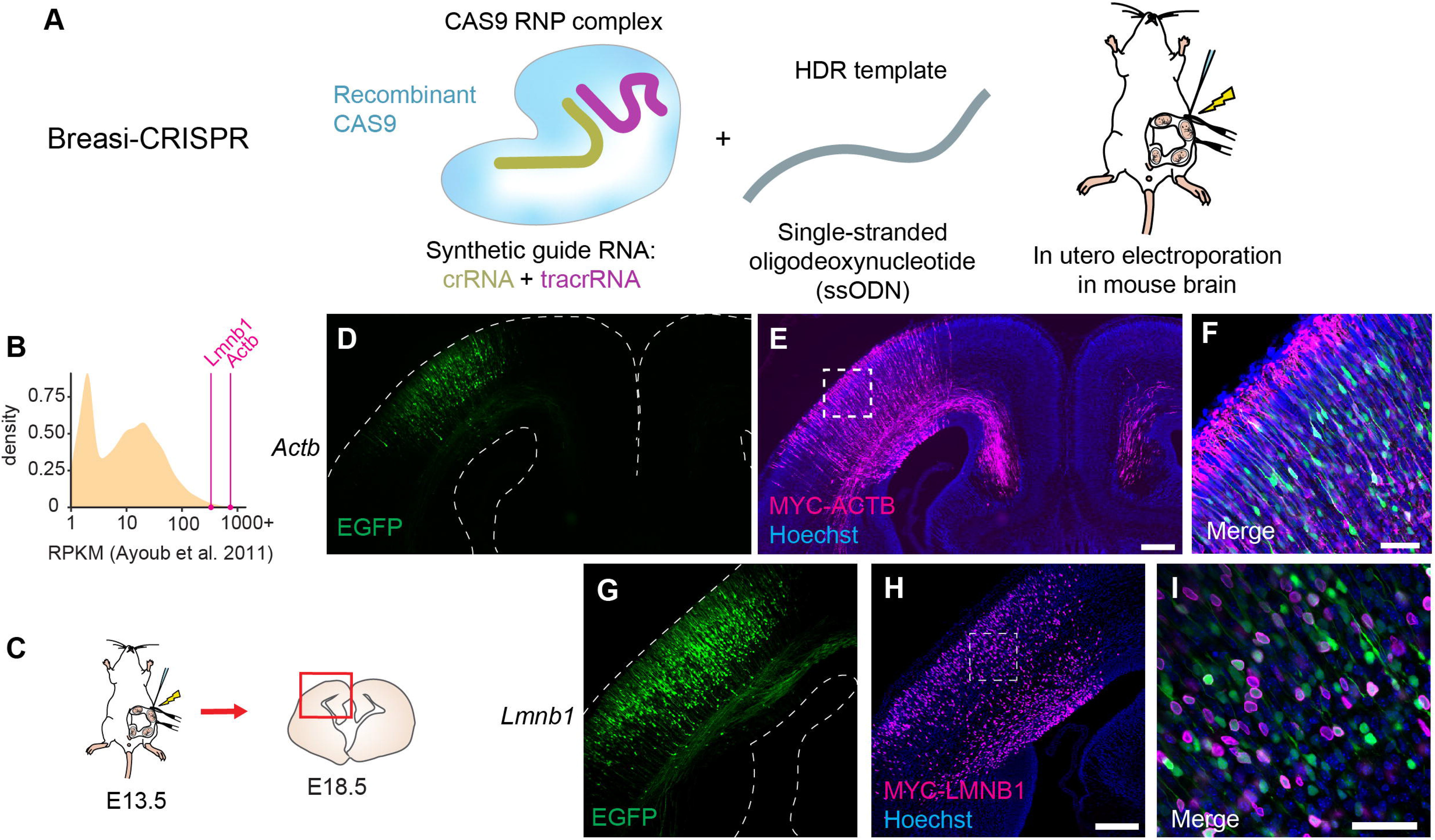
Examples of Breasi-CRISPR results five days after electroporation. (A) Cartoon representing Breasi-CRISPR constituents. (B) Graph showing relative reads per kilobase of exon model per million mapped reads (RPKM) for Lmnb1 and Actb (data from Ayoub et al. 2011). (C) Cartoon outlining approach for Fig. 1 D-I. Breasi-CRISPR IUE was performed at E13.5 followed by sample collection at E18.5 and histological analysis of coronal sections. (D-F) Representative confocal images of Breasi-CRISPR-tagged MYC-ACTB (fuchsia) in electroporated neurons (green). (G-I) Representative confocal images of Breasi-CRISPR-tagged MYC-LMNB1 (fuchsia) in electroporated neurons (green). Images are representative of >3 brains across several litters. Scale bars: 200 μm (D, G) and 50 μm (E, H).

## Results

### Breasi-CRISPR is an efficient method to epitope-tag various endogenous proteins in the embryonic mouse cortex

As a first test of the efficacy of this technique, we used Breasi-CRISPR to epitope-tag the abundant, ubiquitous proteins BETA-ACTIN and LAMIN-B1 (encoded by the *Actb* and *Lmnb1* genes, respectively). Of note, we used published RNA-seq data from microdissected E14.5 mouse cortices as a proxy for overall protein abundance at the targeted time point (Fig. 1B) (Ayoub et al., 2011). BETA-ACTIN is a cytoskeletal protein important throughout cortical development both in progenitors and neurons (Lian and Sheen, 2015). LAMIN-B1 is a nuclear lamina protein responsible for maintaining nuclear architecture, DNA replication and gene expression (David, 2011). We used single stranded oligodeoxynucleotide (ssODN) HDR templates to integrate the MYC tag sequence immediately downstream of the translation start site of *Actb* and *Lmnb1* genes. IUEs were performed at E13.5, together with a plasmid expressing EGFP to visualize transfected cells (Fig. 1C). Our initial analysis of Breasi-CRISPR targeted cortices was performed at E18.5, five days after the electroporation, since we hypothesized that it would take time for CRISPR reagents to introduce the MYC sequence into the targeted loci, and for cells to express the tagged endogenous proteins. For both gene targets, IHC targeting the MYC tag consistently showed a very large number of cells with positive signal. As expected, MYC-ACTB signal showed filamentous structures present in the cytoplasm, and was excluded from nuclei but prominent in axons crossing the midline (Figs. 1D-F). MYC-LMNB1 was observed along the edge of nuclei, consistent with its role as a component of the nuclear lamina (Figs. 1G-I).

These results prompted us to measure Breasi-CRISPR efficiency within a shorter timeframe following IUE. Therefore, we analyzed signal for MYC-ACTB and MYC-LMNB1 via IHC just two days after electroporation at E13.5 (Fig. 2A). Under these conditions, strong MYC-ACTB signal was observed in numerous radial glial cells in the ventricular zone (VZ), as well as in migrating neurons in the subventricular zone (SVZ) and in the intermediate zone (IZ, Figs. 2C-G). In the IZ and in the cortical plate, MYC-ACTB also highlighted radial glial basal processes and endfeet with little to no cytoplasmic EGFP signal (Figs. 2D,G). The lack of EGFP in those structures is a result of low diffusion in thin processes, and demonstrates that tagging endogenous proteins populating distal subcellular compartments may be advantageous to visualize cellular morphology when diffusion of fluorescent proteins is limited.

**Figure 2.**
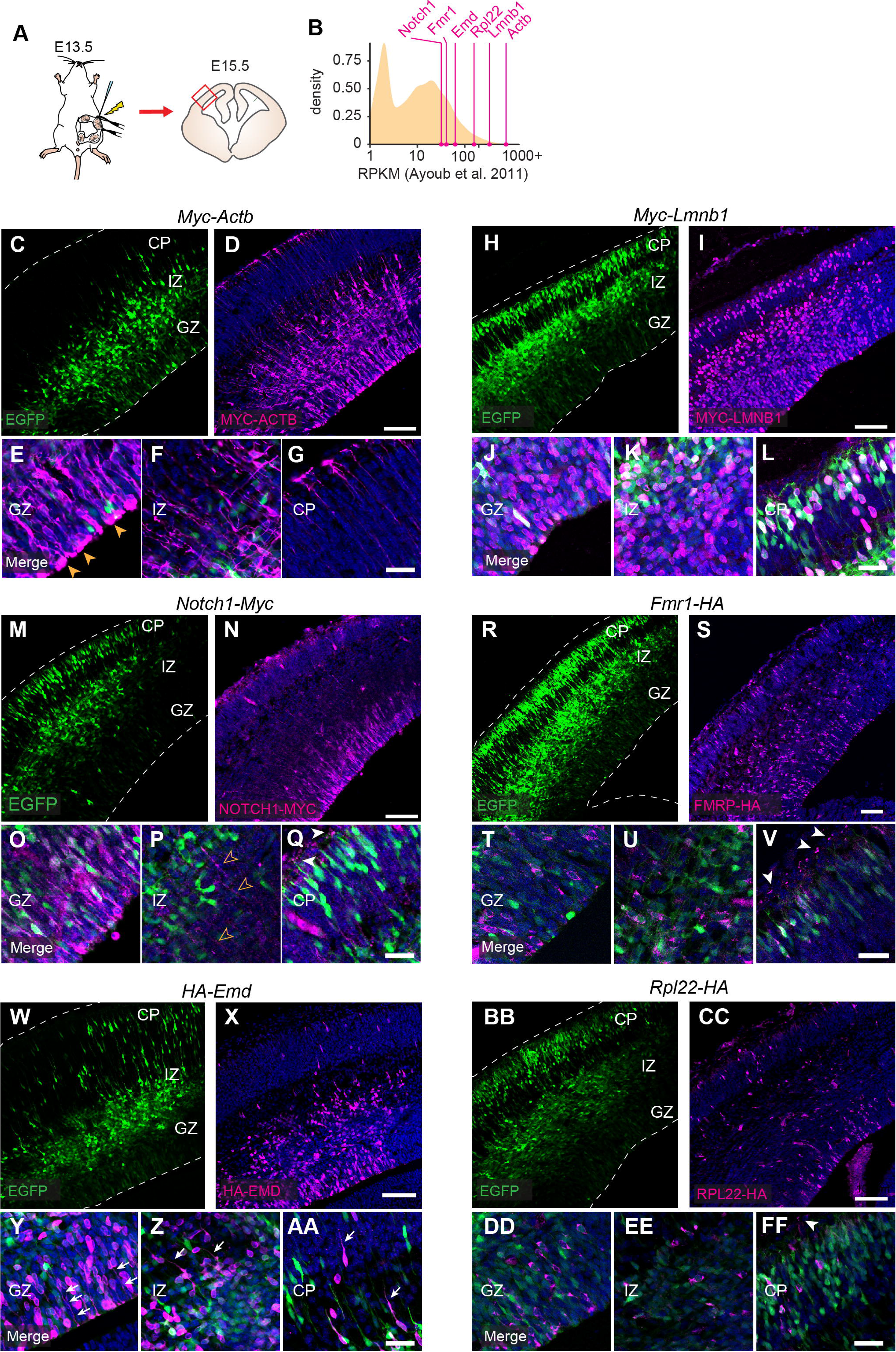
Examples of Breasi-CRISPR results two days after electroporation. (A) Cartoon outlining approach for Fig. 2 C-FF. Breasi-CRISPR IUE was performed at E13.5 followed by sample collection at E15.5 and histological analysis of coronal sections. (B) Graph showing relative reads per kilobase of exon model per million mapped reads (RPKM) for Notch1, Fmr1, Emd, Rpl22, Lmnb1, and Actb (data from Ayoub et al. 2001). (C-G) Representative confocal images of Breasi-CRISPR-tagged MYC-ACTB (fuchsia) in electroporated neurons (green). Yellow arrows (E) point to radial glia actin rings at the ventricular border (E) (H-L) Representative confocal images of Breasi-CRISPR-tagged MYC-LMNB1 (fuchsia) in electroporated neurons (green). (M-Q) Representative confocal images of Breasi-CRISPR-tagged NOTCH1-MYC (fuchsia) in electroporated neurons (green). Yellow arrows (P) and White arrows (Q) point to NOTCH1-MYC signal in radial glia basal processes and endfeet, respectively. (R-V) Representative confocal images of Breasi-CRISPR-tagged FMRP-HA (fuchsia) in electroporated neurons (green). White arrows (V) point to FMRP-HA signal in radial glia basal endfeet.(W-AA) Representative confocal images of Breasi-CRISPR-tagged HA-EMD (fuchsia) in electroporated neurons (green). White arrows (Y, Z, AA) point to unexpected HA-EMD cytoplasmic signal. (BB-FF) Representative confocal images of Breasi-CRISPR-tagged RPL22-HA (fuchsia) in electroporated neurons (green). White arrows (FF) point to RPL22-HA signal in radial glia basal endfeet. Scale bars: 100 μm (D, I, N, S, X, CC) and 30 μm (G, L, Q, V, AA, FF).CP: cortical plate; IZ: intermediate zone; GZ: germinal zone

While these data demonstrate efficacy of Breasi-cRISPR for highly expressed proteins, such as LAMIN-B1 and BETA-ACTN, it is unclear how well it works for lower expressed proteins. To demonstrate that Breasi-CRISPR works not only with highly expressed proteins we targeted four genes with lower reported expression levels (Fig. 2B). We chose *Notch1, Fmr1, Emd*, and *Rpl22*, encoding a transmembrane receptor, an RNA-binding protein, another nuclear lamina protein and a ribosomal protein, respectively. In most cases, we selected where to insert tags within proteins based on previously published data for the protein of interest.

Full-length NOTCH1 is a transmembrane receptor protein involved in the regulation of neural progenitor proliferation (Yoon and Gaiano, 2005). Upon binding to its ligand, the C-terminus of NOTCH1 is cleaved and transported to the nucleus where it regulates transcription of its target genes. In an attempt to capture both the full length (membrane-bound) and cleaved version of NOTCH1 (cytoplasmic and nuclear) in neural progenitors, we inserted the MYC sequence in the portion of the *Notch1 gene* encoding the C-terminus of the protein. Two days after IUE, we observed NOTCH1-MYC signal specifically in neural progenitors, with most of the signal showing membrane localization (Figs. 2M-Q). While we expected to observe more NOTCH1-MYC signal in progenitor nuclei, this could be explained by the known short half-life of the intracellular domain–– compared to that of the full-length protein at the membrane (Carrieri and Dale, 2016). Of note, we observed NOTCH1-MYC signal not only in radial glia soma and apical endfeet, but also in their basal processes and endfeet (Figs. 2N,P,Q). This suggests that NOTCH1 may have an underexplored role in those structures.

Next, we inserted an HA-tag sequence immediately upstream of the stop codon of *Fmr1*, the gene encoding FMRP. FMRP is an RNA-binding protein that controls diverse aspects of mRNA metabolism such as translation and localization. It has been reported to be ubiquitously expressed in the developing cortex with a specific enrichment in radial glia apical and basal endfeet (Saffary and Xie, 2011, Pilaz et al., 2016a). As expected, FMRP-HA was mainly observed in the cytoplasm of targeted cells (Figs. 2R-V). In radial glia, strong signal was observed in apical endfeet as well as in basal endfeet (Figs. 2S, V), as observed previously (Saffary and Xie, 2011, Pilaz et al., 2016a).

*Emd* encodes EMERIN, a nuclear membrane protein and a component of the nuclear lamina (Koch and Holaska, 2014). We used Breasi-CRISPR to insert an HA-tag sequence in the N-terminus of EMERIN. Two days after electroporation, we observed many cells displaying IHC signal for HA resembling the signal we observed for MYC-LMNB1 (Figs. 2W-AA). This suggested that the HA tag sequence was correctly inserted in the *Emd* locus. However, we also observed many cells with localized cytosolic IHC signal. Some radial glia and migrating neurons showed localization in their apical and leading process, respectively (Figs. 2X-AA). Since localization of EMERIN has been reported outside of the nucleus at adherens junctions in cardiomyocytes (Wheeler et al., 2010), it is tempting to speculate that the signal we observed suggests a previously unknown extra-nuclear role for EMERIN in the developing cortex.

We also successfully added an HA-tag to the C-terminus of RPL22 via Breasi-CRISPR. We used this strategy with this specific protein, since its endogenous C-terminal tagging has been used to generate the Ribo-tag mouse line (Sanz et al., 2009), enabling the pull down of ribosome-bound RNAs in specific cell types. After two days, RPL22-HA signal was cytoplasmic and observed in radial glia as well as migrating neurons (Figs. 2BB-FF). Of note, we also observed RPL22-HA signal in the radial glia basal endfeet region within the marginal zone (MZ, Fig. 2FF). The localization of the RPL22-HA signal is in line with the known role of RPL22 and ribosomes in the developing mouse cortex.

Finally, we tested whether CRISPR-tagging could be detected as soon as 24 hours after IUE (Fig. 3A). This was performed via Breasi-CRISPR tagging of ACTB or LMNB1, introducing the MYC sequence immediately after the start codons of *Actb* and *Lmnb1*, respectively. We were surprised to observe a significant number of cells displaying MYC signal for both proteins (Figs. 3B-E), suggesting that the genomic integration of the targeted sequence can happen very rapidly after IUE. Altogether, these experiments highlight that Breasi-CRISPR can be used to tag a variety of endogenous proteins expressed during embryonic development. This will be particularly useful when proteins of interest lack efficient targeting antibodies. This also permits the visualization of those proteins in isolated cells and, thus, a more reliable assessment of protein localization within those cells, which can be difficult when proteins of all the cells within a tissue are revealed by IHC.

**Figure 3.**
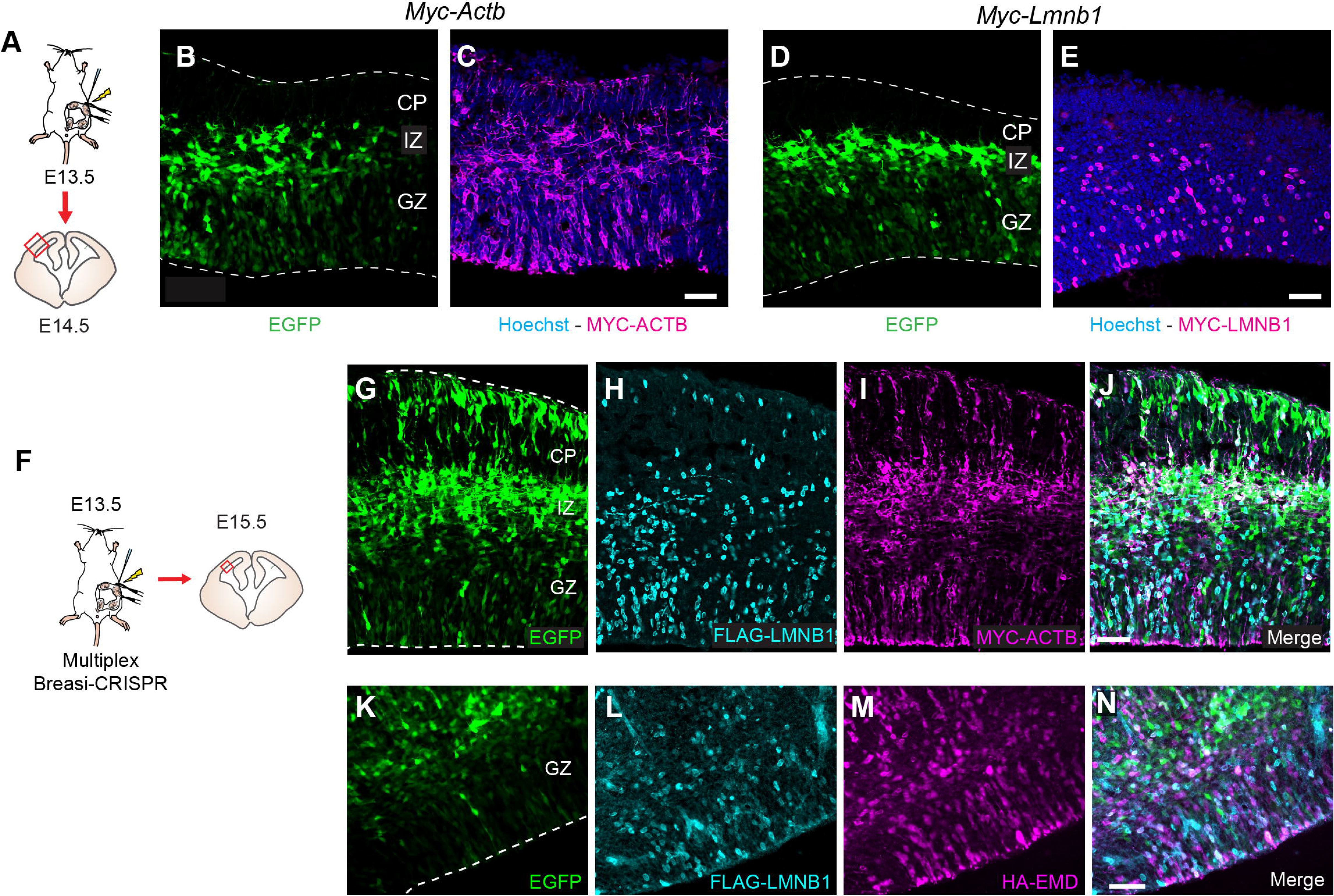
Examples of Breasi-CRISPR results one day after electroporation and multiplexing. (A) Cartoon outlining approach for Fig. 3 B-E. Breasi-CRISPR IUE was performed at E13.5 followed by sample collection at E14.5 and histological analysis of coronal sections. (B-C) Representative confocal images of Breasi-CRISPR-tagged MYC-ACTB (fuchsia) in electroporated neurons (green). (D-E) Representative confocal images of Breasi-CRISPR-tagged MYC-LMNB1 (fuchsia) in electroporated neurons (green). (F) Cartoon outlining approach for Fig. 3G-N. Breasi-CRISPR IUE was performed at E13.5 followed by sample collection at E15.5 and histological analysis of coronal sections. (G-J) Representative confocal images of multiplex Breasi-CRISPR-tagged FLAG-LMNB1 (turquoise) and MYC-ACTB (fuchsia) in electroporated neurons (green). (K-N) Representative confocal images of multiplex Breasi-CRISPR-tagged FLAG-LMNB1 (turquoise) and HA-EMD (fuchsia) in electroporated neurons (green). Scale bars: 50 μm (C-E, G-N).CP: cortical plate; IZ: intermediate zone; GZ: germinal zone.

### Multiplexing Breasi-CRISPR

We multiplexed the Breasi-CRISPR approach in order to tag multiple different proteins in the same brain region at the same time. First, we targeted *Lmnb1* and *Actb* simultaneously, adding FLAG- and MYC-tags to LMNB1 and ACTB, respectively (Figs. 3F-J). Next, we targeted LMNB1 and EMERIN, both proteins of the inner nuclear membrane (Figs. 3K-N). For each, we used two crRNAs: one crRNA targeting the *Lmnb1* gene and the other crRNA targeting the *Emd* or the *Actb* genes. Together with these crRNAs, we electroporated two ssODNs: one to add a FLAG-tag in the *Lmnb1* gene and one to add a MYC- or HA-tag in the *Actb* or *Emd* gene respectively. We performed IUEs at E13.5 and collected the samples at E15.5. IHC analyses show abundant tagging of both proteins, mostly in the same cells (Figs. 3G-N). This demonstrates that Breasi-CRISPR can be used to study the co-localization of multiple proteins within individual cells *in vivo*.

### Quantification of Breasi-CRISPR efficiency

To quantify Breasi-CRISPR efficiency, we first used MYC-tagging of LAMINB1 since this protein is ubiquitously expressed and its localization in the nucleus facilitates cell counting. We performed quantifications two days and five days after IUE at E13.5 (Figs. 2A-J). Within the electroporated region, we counted 43% EGFP+ cells amongst Hoechst+ nuclei, and within these EGFP+ cells, 42% showed MYC signal. However, 48% of the MYC+ nuclei were not EGFP+ (Figs. 2 B-E). After five days, we observed that 17% of Hoechst+ nuclei were EGFP+, and less than half of those EGFP+ nuclei were MYC+. 72% of the MYC+ nuclei were not EGFP+ (Figs. 2F-J). The large fraction of MYC+, GFP-negative cells was expected because while the genomic integration of the MYC sequence is permanent and inherited by all the progeny of targeted progenitors, the cycling progeny of transfected cells lose EGFP expression due to the dilution of the plasmid as they progress through multiple cell cycles. Therefore, based on Hoechst counterstaining, we also quantified the total fraction of nuclei with efficient Breasi-CRISPR-mediated tagging. We observed that 34% and 27% of all the nuclei were positive for MYC signal after two days and five days, respectively (Figs. 4E,J). Altogether, this data shows that Breasi-CRISPR tagging of proteins in the developing cortex is extremely efficient.

**Figure 4.**
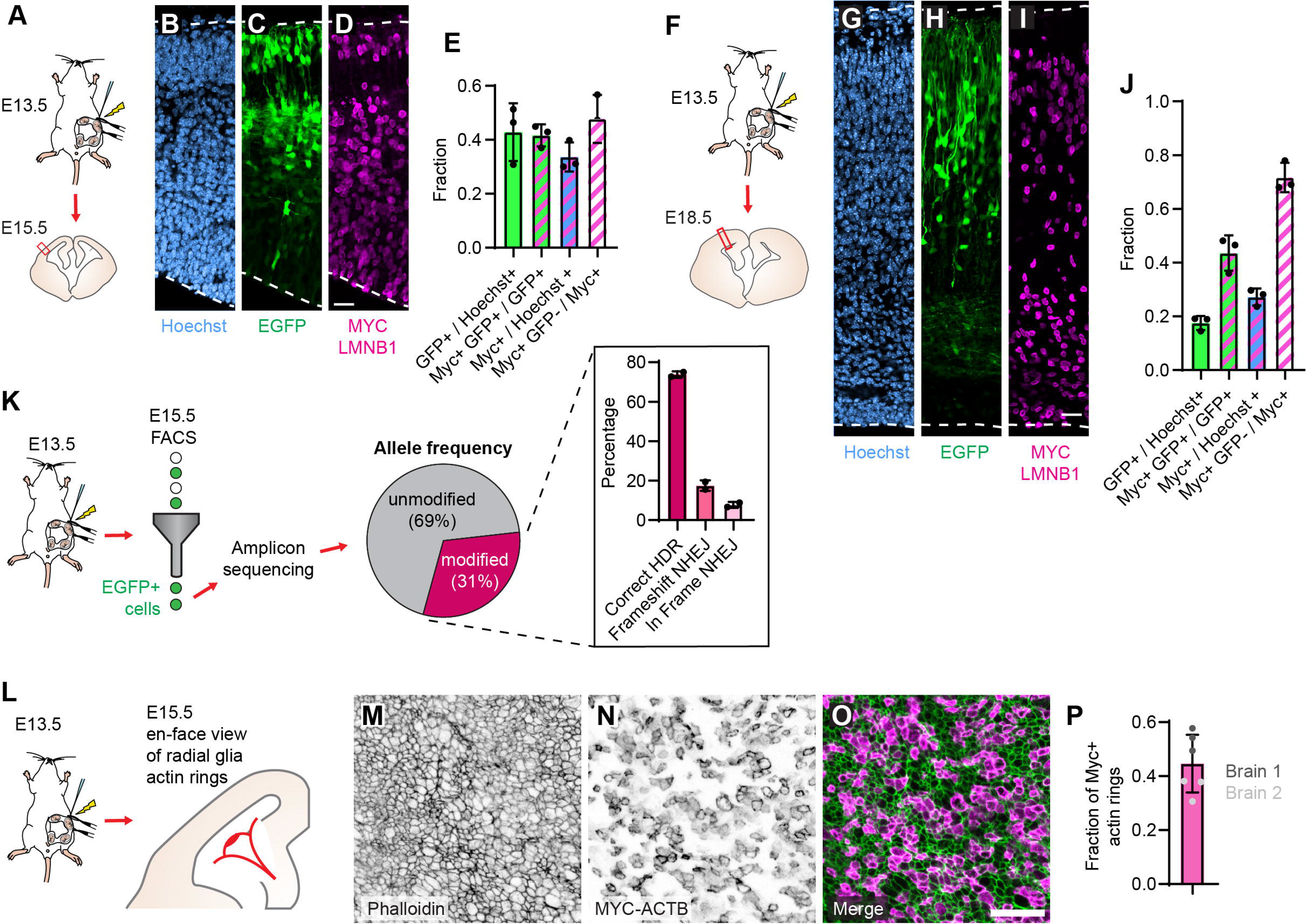
Quantification of Breasi-CRIPSR efficiency. (A) Cartoon outlining approach for Fig. 4 B-E. Breasi-CRISPR IUE was performed at E13.5 followed by sample collection at E15.5 and histological analysis of coronal sections. (B-D) Representative confocal images of Breasi-CRISPR-tagged MYC-LMNB1 (fuchsia) in electroporated neurons (green) with Hoescht (blue). (E) Quantification of Breasi-CRISPR efficiency via cell counting two days after Breasi-CRISPR IUE (n=3 brains from two different liters). (F) Cartoon outlining approach for Figs. 4 G-J. Breasi-CRISPR IUE was performed at E13.5 followed by sample collection at E18.5 and histological analysis of coronal sections. (G-I) Representative confocal images of Breasi-CRISPR-tagged MYC-LMNB1 (fuchsia) in electroporated neurons (green) with Hoescht (blue). (J) Quantification of Breasi-CRISPR efficiency via cell counting five days after Breasi-CRISPR IUE (n=3 brains from two different litters). (K) Cartoon outlining approach for sequencing validation. Breasi-CRISPR IUE was performed at E13.5 followed by sample collection at E15.5 with FACS sorting for GFP+ cells and amplicon sequencing. Percent of cells with correct HDR, frameshift NHEJ, or in frame NHEJ were quantified (n=2 litters). (L) Cartoon outlining approach for Figs. 4N-Q. Breasi-CRISPR IUE was performed at E13.5 followed by sample collection and phalloidin staining of wholemount cortices. (M-P) Representative confocal images of Breasi-CRIPSR-tagged MYC-ACTB (fuchsia) stained with phalloidin (F-actin stain, green). (P) Quantification of Breasi-CRISPR efficiency via quantification of MYC+ actin rings (n=2 brains). Scale bars: 25 μm (D, J) and 20 μm (P). FACS: Fluorescence-activated cell sorting, HDR: Homology-directed repair, NHEJ: non-homologous end joining

To further examine the efficacy of Breasi-CRISPR, we used amplicon sequencing to quantify the gene editing products generated by the approach using MYC-LMNB1. We performed IUEs at E13.5 and employed fluorescence activated cell-sorting at E15.5 to isolate GFP+ cells. Using DNA extracted from these cells, we amplified the region surrounding the Breasi-CRISPR-induced modification by polymerase chain reaction (PCR). PCR products were purified and subsequently analyzed by next-generation sequencing (Fig. 4K). The sequencing results showed that out of all the *Lmnb1* sequences, 23.4% presented the correct HDR-mediated addition of the MYC tag sequence to the *Lmnb1* gene. Out of all the modified sequences, the percentage of correct HDR modified sequences was greater than 70% (73.8%) and the NHEJ events frequencies were lower than 30% (26.2%), with a fraction of indels leading to frameshifts equal to 17.7% (Fig. 4K). These amplicon sequencing results show that Breasi-CRISPR can efficiently add an epitope-tag to coding sequences in the embryonic cortex within two days of IUEs with minimal generation of indels which could be deleterious for the expression of the gene of interest, and hence the viability of the targeted cells. This makes it ideal for studying proteins that are expressed transiently during development.

As another means to test Breasi-CRISPR efficacy using another gene target, we analyzed MYC-ACTB signal together with the F-ACTIN marker phalloidin in wholemounts of electroporated cortices, focusing on the ventricular border 2 days after IUE (Figs. 4L-P). This allowed us to visualize MYC-ACTB+, Phalloidin+ “actin-rings” localized at radial glia apical endfeet. Measuring the fraction of MYC-ACTB+, Phalloidin+ endfeet within the whole Phalloidin+ population, this approach enabled us to quantify the fraction of radial glia in which BETA-ACTIN was epitope-tagged by Breasi-CRISPR. In the electroporated regions, we counted that 30% to 57% of all Phalloidin-labelled apical endfeet showed MYC-ACTB signal. Altogether this data demonstrates the rapid efficacy of the Breasi-CRISPR approach.

### Live imaging of endogenous proteins fused to fluorescent proteins via Breasi-CRISPR

Next, we performed a set of experiments to show that Breasi-CRISPR allows the knock-in of larger tags such as EGFP, thus enabling live imaging studies of endogenous proteins during cortical development. For this, we used ssODNs with EGFP tag and sequence targeting *Lmnb1* gene. These ssODNs were generated in house using the ivTRT method (Quadros et al., 2017, Miura et al., 2018). First, IUEs were performed at E13.5 and brains were collected and fixed at E18.5, using tdTomato or mCherry expression plasmids as a marker of transfection (Figs. 5A-C). We found 20% of the tdTomato cells showed EGFP-tagged LAMIN-B1 and represented 1% of all the nuclei within the electroporated region. We then visualized Breasi-CRISPR-tagged EGFP-LMNB1 by live imaging in acute embryonic brain slices. Live slices were generated at E15.5, two days after the Breasi-CRISPR IUE (Fig. 5D). Live imaging was performed overnight for 20 hours. With this approach, we were able to visualize the interkinetic nuclear migration of neural stem cell nuclei in the VZ, and the migration of neurons in the SVZ and IZ (Supplementary Movie 1, Figs. 5F). In migrating neurons, tensions applied to nuclei as the cells migrate through dense tissue was evident (Fig. 5F). Altogether, this data demonstrates that Breasi-CRISPR can be utilized to visualize endogenous protein dynamics during embryonic cortical development. This provides a powerful approach which is not reliant on exogenous expression of proteins or transgenic mouse lines.

**Figure 5.**
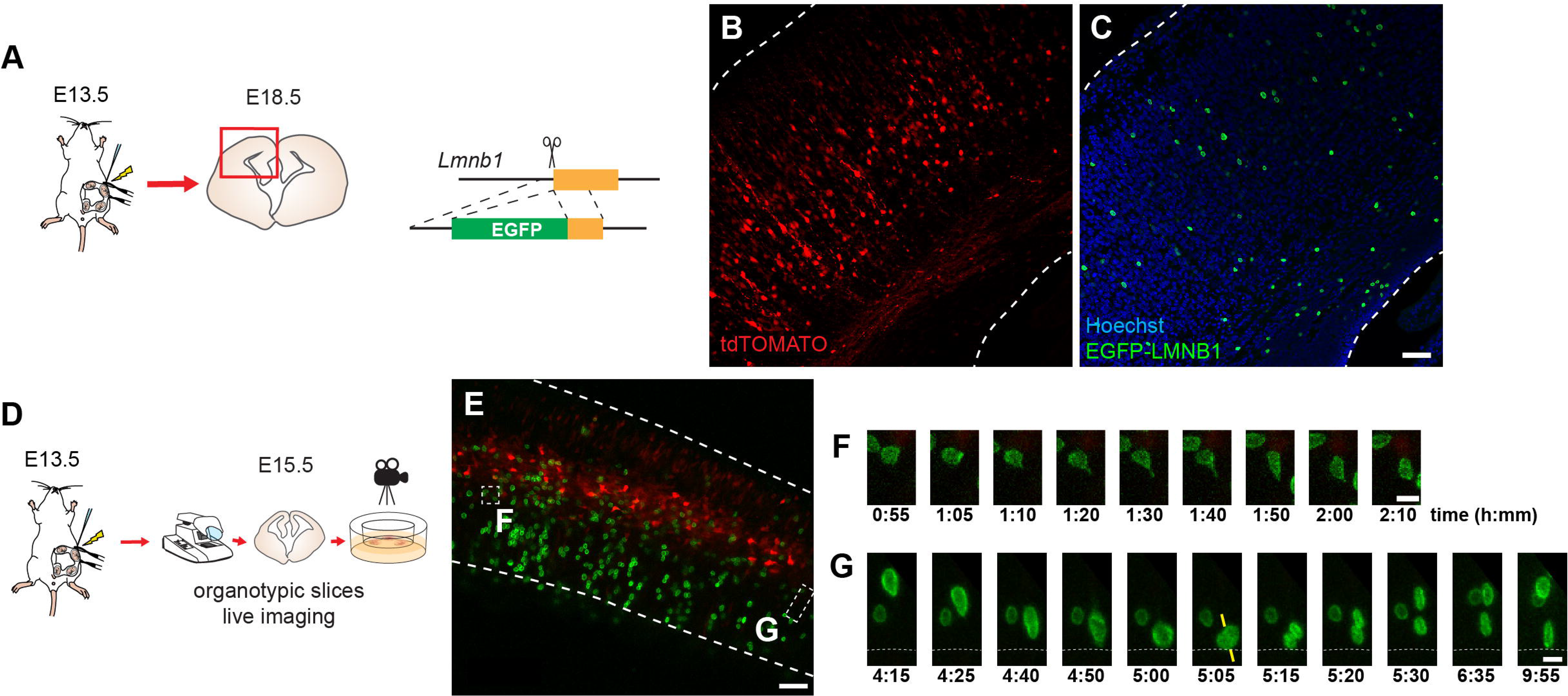
Fluorescent tagging with Breasi-CRISPR. (A) Cartoon outlining approach for B,C. Breasi-CRISPR IUE was performed at E13.5 followed by sample collection at E18.5 and histological analysis of coronal sections. (B,C) Representative confocal images of Breasi-CRISPR-tagged EGFP-LMNB1 (green) in electroporated neurons (red). (D) Cartoon outlining approach for E-G. Breasi-CRISPR IUE was performed at E13.5 followed by sample collection at E15.5, organotypic slice culture, and live imaging. (E) Representative confocal image of Breasi-CRISPR-tagged EGFP-LMNB1 (green) in electroporated neurons (red). (E-G) Representative stills from time-lapse confocal images of Breasi-CRISPR-tagged EGFP-LMNB1 (green). Yellow bars (G) indicate mitotic cleavage plane. Scale bars: 50 μm (B,C,E) and 5μm (F,G)

### Using Breasi-CRISPR to study protein-protein interactions

Given the efficiency we observed by IHC, we were encouraged to test if the Breasi-CRISPR technique would enable us to perform in-depth protein studies using tagged proteins in the developing cortex. First, we attempted to detect tagged proteins using immunoblot analyses of lysates two days after IUE at E13.5 (Fig. 3A). For MYC-ACTIN (Fig. 6B), MYC-LAMIN B1 (Fig. 6D), and HA-EMD (Fig. 6E), we were able to detect the tagged proteins within lysates from one, three and four cortices, respectively. Additionally, we were able to immunoprecipitate MYC-ACTIN, using a MYC antibody (Fig. 6C). This data further demonstrates the efficiency of Breasi-CRISPR, prompting us to attempt to use Breasi-CRISPR for downstream high-throughput proteomics studies such as co-immunoprecipitation (co-IP). Thus, we performed co-IP mass spectrometry analyses to reveal interactors of FMRP, a protein of medium abundance (Fig. 2B) in an unbiased fashion. To do this, we used Breasi-CRISPR to knock in the HA tag immediately upstream of the stop codon of FMRP. We performed IUEs at E13.5 and collected cortex samples at E15.5. Pooled cortices from single litters (2 litters with 5-6 embryos per litter) were utilized as technical replicates for co-IP experiments, and co-IPs of non-electroporated cortices were used as controls (Figs. 7A,B). Consistent with the known role of FMRP in the regulation of mRNA translation, gene ontology analyses of proteins significantly enriched in the co-IP fractions from FMRP-HA cortices showed an over-representation of networks linked to mRNA processing, especially for ribosomes and the spliceosome (Figs. 7C-E, Table S1). We also observed an enrichment of intermediate filament proteins, which are regulated by FMRP (Thomsen et al., 2013). Surprisingly, we also found a network specific to complement activation. We were also initially surprised to discover an enrichment for a network of histone proteins, however direct interactions between FMRP and this subclass of protein have been reported (Alpatov et al., 2014). Altogether, this demonstrates that Breasi-CRISPR can be used to reveal protein-protein interactions in the developing mouse cortex.

**Figure 6.**
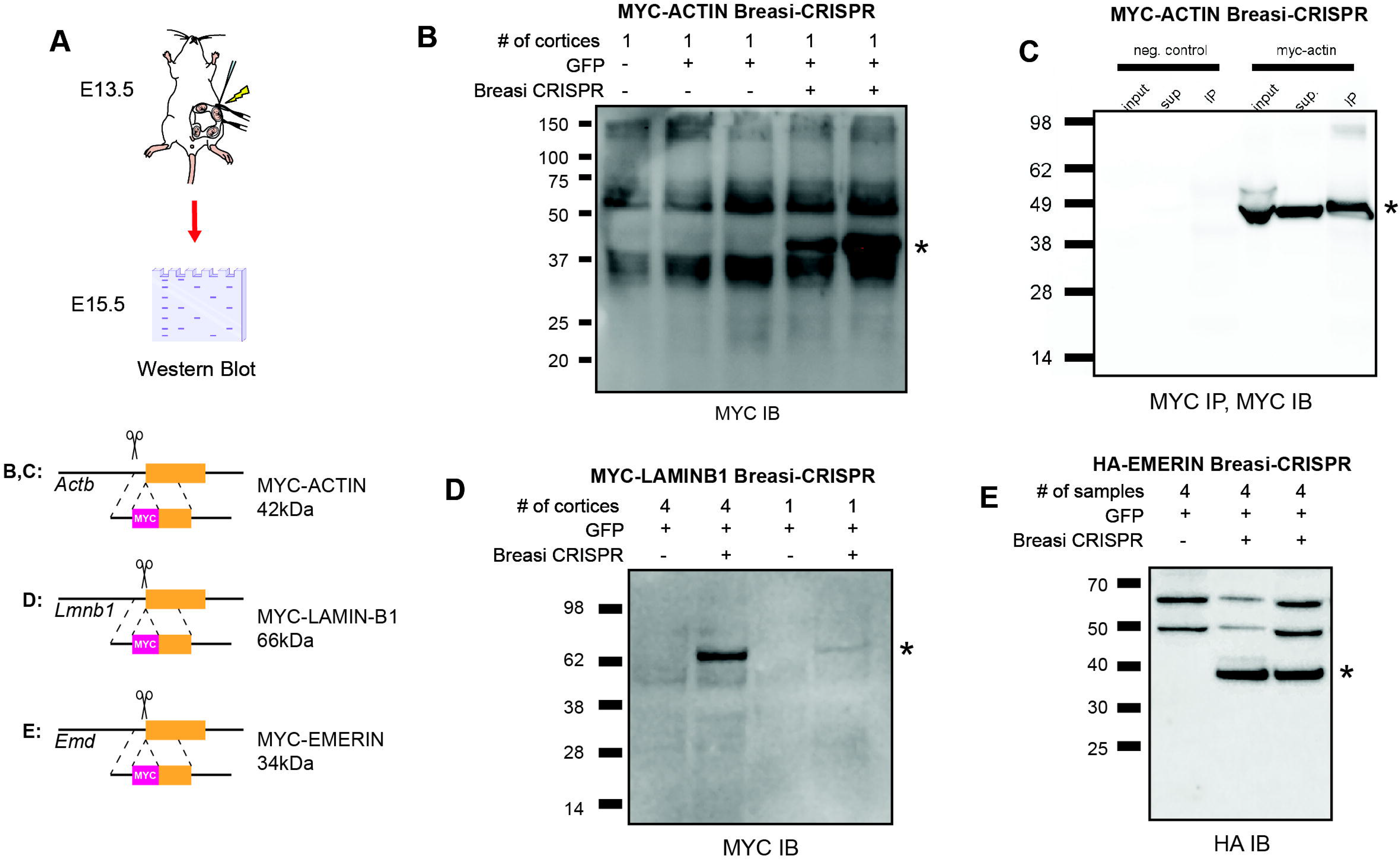
Visualization of Breasi-CRISPR-tagged proteins by immunoblotting. (A) Cartoon outlining approach for Figs. 6 B-E. Breasi-CRISPR IUE was performed at E13.5 and cortices were collected for Immunoblots (IB) at E15.5. Also depicted are cartoons indicating the tag insertion with the molecular weight (in kDa) of the protein and tag together. (B) Representative IB showing increased MYC signal in MYC-ACTIN Breasi-CRISPR electroporated samples compared to samples electroporated with GFP alone. (C) Representative IB showing enrichment of MYC signal in MYC-ACTIN Breasi-CRISPR electroporated samples compared to samples electroporated with GFP alone. (D) Representative IB showing increased MYC signal in MYC-LAMINB1 Breasi-CRISPR electroporated samples compared to samples electroporated with GFP alone, and an increase when four cortices are pooled compared to one cortex alone. (E) Representative IB showing increased HA signal in HA-EMERIN Breasi-CRISPR electroporated samples compared to samples electroporated with GFP alone. IP: immunoprecipitation; IB: immunoblot.

**Figure 7.**
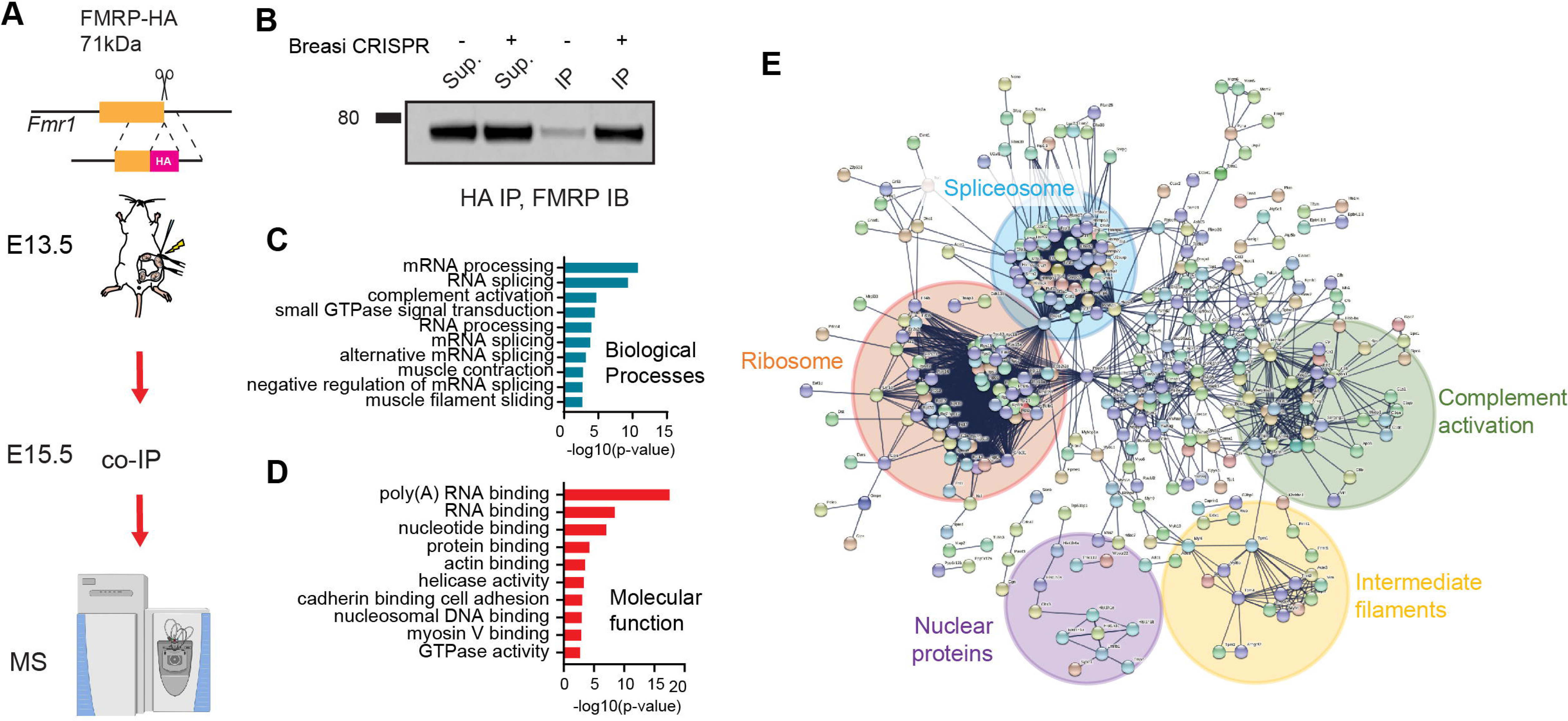
Example of Co-IP/MS with Breasi-CRISPR. (A) Cartoon outlining approach for Figs. 7 B-E. FMRP-HA Breasi CRISPR IUE was performed at E13.5 and cortices were collected for Co-IP/MS at E15.5. (B) Representative IB showing enrichment of FMRP signal following HA IP in FMRP-HA electroporated samples compared to samples electroporated with GFP alone. (C-D) Gene ontology analyses showing biological processes (C) and molecular functions (D) enriched in FMRP-HA Breasi-CRISPR electroporated samples. (E) Network analysis of genes enriched in FMRP-HA Breasi-CRISPR from MS analysis. IP: immunoprecipitation; IB: immunoblot; MS: mass spectrometry

## Discussion

In this study we describe Breasi-CRISPR, an improved approach combining IUE with CRISPR technology to tag endogenous proteins in the developing mouse cortex. Breasi-CRISPR enabled us to tag proteins in up to 30% of all the cells within the electroporated area. Tagged proteins can be visualized by IHC as soon as one day following electroporation and by immunoblotting as soon as two days after IUE. Breasi-CRISPR also shows multiplexing capacity and enables the insertion of fluorescent tags to visualize the dynamics of endogenous protein in large numbers of cells, via imaging in live tissue. Moreover, using tagged FMRP as a bait, we demonstrate that Breasi-CRISPR can be utilized to interrogate protein-protein interactions by co-immunoprecipitation of tagged proteins. Current methods for these approaches rely upon effective antibodies, which may not be available for a protein of interest, or on overexpression of tagged proteins, which can lead to non-specific binding. This technique will propel an understanding of cortical development by allowing specific detection under endogenous expression by overcoming those challenges.

Published approaches coupling CRISPR technology with IUE to tag endogenous proteins rely on plasmids expressing the CAS9 protein and the gRNA (Tsunekawa et al., 2016, Suzuki et al., Mikuni et al., 2016, Uemura et al., 2016, Fang et al., 2021). Thus, there is a significant lag between electroporation and the production of CAS9 within electroporated cells, emanating from the need for the plasmid to integrate the nucleus, followed by transcription of the mRNA encoding CAS9, the export of this mRNA to the cytoplasm and finally its translation. Since integration of the plasmid into the nucleus requires the cells to accomplish mitosis soon after IUE (Stancik et al., 2010, Pilaz et al., 2009), using plasmids encoding CRISPR reagents limits the number of cells eventually expressing CAS9 and the gRNA. With the Breasi-CRISPR approach however, a pre-formed CAS9/gRNA complex is directly delivered into the cells and is capable of targeting the genome immediately. Moreover, since the gRNAs are synthetic, they can be chemically enhanced to promote their stability, therefore maintaining high levels in targeted cells for longer periods of time. This is also true for single-stranded HDR templates. However, due to cost, longer ssODN HDR templates introducing fluorescent tags are usually generated in-house through ivTRT (Quadros et al., 2017). In this case, to our knowledge adding stabilizing chemical modifications to the ssODN is not possible, although this could be a valuable avenue of research in the future.

Like any other CRISPR-based approaches, Breasi-CRISPR may have off-target effects in electroporated cells. This can include undesirable editing of other genomic loci due to sequence-homology with the targeted sequence. Because of the error-prone nature of the DNA-repair pathways, this could lead to frameshifts when coding-regions are affected, or alteration of critical regulatory regions when non-coding regions are disrupted. Many online algorithms allow the experimenter to predict gRNA off-target sequences, enabling the mitigation at the design level. However, undesirable editing could also occur in the targeted region itself due to potential inefficiency of the HDR-based approach. We tested this directly through sequencing of the targeted region and observed NHEJ-induced frameshifts in 5% of all the sequenced fragments (compared with 23% for correct HDR), suggesting that, overall, this effect may be minimal. Additionally, we did not see any increase of apoptosis in electroporated brains, and the distribution of EGFP+ cells did not seem altered (comparing brains targeted with and without Breasi-CRISPR, data not shown). This suggests that Breasi-CRISPR does not have overt noxious effects.

Most of our Breasi-CRISPR designs worked efficiently without the need for optimization. However, some required us to test different conditions to find tags and positions within the proteins to enable their visualization by IHC or immunoblotting. We assumed that this is because certain epitope-tags can lead to instability of the protein (Saiz-Baggetto et al., 2017). Additionally, sequences surrounding the epitope-tag can hinder their accessibility to antibodies (Schuchner et al., 2020). One way to mitigate this issue is to add a linker sequence between the tag and the protein (Chen et al., 2013). Moreover, the target protein may be cleaved, leaving the epitope-tag subject to degradation. Finally, the target protein may be expressed at levels so low that its detection might be problematic. In that case, it is possible to add tandem-repeats of the tag instead of single copies. Of note however, the efficiency of the integration might be affected since it is inversely correlated with the size of the inserted sequence (Ohtsuka et al., 2018).

Breasi-CRISPR is expected to work best at earlier developmental time points when progenitors preferentially use HDR over NHEJ for DNA repair (Mikuni et al., 2016). In this study, we performed all the Breasi-CRISPR experiments at E13.5, which is the earliest time at which our group is reproducibly efficient at IUE (75% average success rate). However, the efficacy of the Breasi-CRISPR approach may be further improved when performed at even earlier time points, but will probably decrease at later timepoints as observed for the SLENDR approach. This limits the opportunity to specifically target cells born towards the end of corticogenesis, such as upper layer neurons and astrocytes. Another consideration related to the timing of experiments is the age at which electroporated brains are collected. While we restricted our collections to embryonic stages, there will be great value in utilizing Breasi-CRISPR to analyze protein localization and interactions in more mature cells during post-natal stages.

In addition to interrogating protein-protein interactions, Breasi-CRISPR should be compatible with RNA-immunoprecipitation to discover the RNA targets of RNA-binding proteins, or with CUT&RUN to shed light on transcription factor binding regions in the genome. It may also be possible to couple the protein-tagging capacity of Breasi-CRISPR with other approaches affecting gene expression, such as RNA interference, or overexpression to test how protein localization and function may be affected by altered expression of other genes. Additionally, multiplexing Breasi-CRISPR for single targets may be possible, enabling the introduction of an epitope tag and a mutation of interest. For instance, this would enable researchers to assess how mutations of certain genes affect the localization of their encoded proteins. Finally, this approach should be applicable to organoids since electroporation-mediated transfection is a viable approach in this system (Lancaster et al., 2013). Together, we anticipate that the multiple applications of Breasi-CRISPR will be a powerful approach enabling researchers to interrogate basic mechanisms related to brain development and function.

## Materials and Methods

### Animals

All experiments were performed in accordance with the Sanford Research IACUC guidelines. C57BL/6J mice were purchased from the Jackson Laboratory and maintained as a breeding colony. Plug dates were defined as embryonic day (E) 0.5 on the morning the plug was identified. To collect the embryonic brain samples, pregnant females were euthanized by CO_2_ inhalation followed by cervical dislocation. Embryonic brains were rapidly collected and evaluated for fluorophore expression under an epifluorescence dissecting microscope (Nikon SMZ3000 stereomicroscope).

### Immunohistochemistry

For immunohistochemistry (IHC) studies, the embryonic brains were fixed with freshly prepared 4% paraformaldehyde (PFA) in PBS overnight, cyro-preserved in 30% sucrose in Phosphate-Buffered Saline (PBS), and frozen in Optimum Cutting Temperature (OCT) compound. Brains were cryo-sectioned at 20μm and attached to charged slides. For IHC, sections were permeabilized using 0.5% Triton X-100 in PBS for 20 min and washed in PBS. The sections were blocked using 10% normal goat serum (NGS) and 1% BSA in PBS for an hour, washed with PBS, and incubated in primary antibody overnight at 4° C. Following PBS washes, sections were incubated with an Alexa Fluor-conjugated secondary antibody and Hoescht for an hour at room temperature (RT), washed in PBS, and mounted in aqueous mounting medium. Antibodies used in this study are indicated in Table S1. IHC-treated sections were imaged on a Nikon A1plus inverted confocal microscope using 10x, 20x, 40x and 60x objectives.

### Amplicon Sequencing

For Amplicon sequencing studies, GFP-expressing cortical regions were dissected under an epifluorescence dissecting microscope. Cells were dissociated via cold protease digestion. The cells were FACS-sorted to collect only GFP-expressing cells. Next, cells were lysed, and genomic DNA was collected. PCR was used to amplify the targeted region in the *Lmnb1* gene. Next, adaptor sequences specific to our Illumina sequencing approach were added via PCR. The PCR product was purified using AxyPrep MagTM PCR Clean-up method. The purity of the eluted DNA was validated using the Bioanalyzer, and the samples were sent to the Sanford Burhnam Genomics Core Facility for Amplicon sequencing.

### Immunoprecipitation

For Immunoprecipitation (IP) studies, GFP-expressing cortical regions were dissected under an epifluorescence dissecting microscope. The isolated tissue was lysed using NP-40 lysis buffer (50 mM Tris-HCl, pH 7.4 at 4°C, 150 mM NaCl, 1 mM EDTA; supplemented with 0.5% NP40). Samples were centrifuged at 12000rpm for 15min, and the supernatant was collected for IP. Magnetic beads with Myc or HA antibodies were used to capture Myc- or HA-tagged proteins from the samples by mixing the beads with the samples for 2-3 hrs at 4 °C. The beads bound to the Myc- or HA-tagged proteins were captured using a magnetic stand and washed with TBST before collection into sample buffer for immunoblotting or freezing for mass-spec analysis. Positive hits were identified based on significant enrichment of peptides in the Breasi-CRISPR electroporated samples compared to non-electroporated samples. There were two replicates per condition. Enrichment was deemed significant when peptides were observed only in Breasi-CRISPR electroporated samples or when the p-value for a t-test comparing the label-free quantitation (LFQ) intensities between eledctroporated and non-electroporated samples was inferior to 0.05, and the ratio between the LFQ intensities between eledctroporated and non-eledctroporated samples was superior to 1.

### Immunoblot analyses

Samples lysed with NP-40 or RIPA buffer were loaded on 4-12% Bolt bis-tris gels and subjected to electrophoresis for 1hr at 200V. The proteins were transferred to nitrocellulose membranes using the Power Blotter system, blocked with 5% milk for 1hr, and incubated with primary antibody overnight at 4° C. Blots were washed with TBST, incubated with HRP-conjugated secondary antibody for an hour at room temperature, washed in TBST, and imaged via chemiluminescence. Antibodies used for the immunoblot analyses are indicated in Table S1.

### *In utero* electroporation (IUE)

IUEs were performed as previously described (Pilaz et al., 2016a). Briefly, E13.5 pregnant mice were anesthetized using isoflurane. An incision was made in the medial ventral abdomen of the pregnant mouse to expose the uterine horns. The Breasi-CRISPR solutions were injected into the lateral ventricles of the embryos and electroporated using tweezer electrodes with the following settings: 60V, 5 square pulses, 60ms duration, 1s intervals. Following the IUE, the uterine horns were returned to the abdominal cavity, the incision was sutured, and the mouse was allowed to recover on a heating pad.

### Preparation of Breasi-CRISPR reagents

The design of the crRNA and ssODN combinations was performed using the Alt-R HDR Design tool on the IDT website. Preparation of Breasi-CRISPR reagents for IUEs followed the protocol established for the iGONAD protocol (Gurumurthy et al., 2019). Briefly, oligonucliotides and CAS9 were ordered from Integrated DNA Technologies (IDT) using the Alt-R design tools and reagents. This includes a transactivating CRISPR RNA (tracrRNA) (IDT #1072532) and targeted CRISPR RNA (crRNA) (sequences and catalog numbers in table S1). The tracrRNA and crRNA (400 μM stock concentration for both) were hybridized in a 1:1 molar solution at 98 °C for 2 minutes then held at room temperature for 10 minutes, resulting in the final guide RNA (gRNA). The final Breasi-CRISPR solution was composed of: 1.5μl of gRNA solution (200μM stock concentration), 2μl HDR template (200μM stock solution for synthetic ssODN, 1-3.5μg/μl for template prepared in house), 1μl CAS9 nuclease (IDT #1081058, stock concentration 10mg/ml), pCAG-GFP/tdTomato/mCherry (0.6μg/μl final concentration), 1μl Fast Green, and RNAase-, DNAase-free water to a final volume of 10μl. This injection solution was incubated at 37° C for 10-30 minutes prior to the surgery. Of note, inserts used for EGFP-LMNB1 Breasi-CRISPR experiments were generated in-house using the ivTrt approach (Quadros et al., 2017). PCR primers used to generate the template for in vitro transcription can be found in Table S1.

### Preparation of live brain slice and live imaging

Brain slices for live imaging were prepared as described previously (Pilaz and Silver, 2014, Pilaz et al., 2016b). Briefly, brains were collected two days post-IUE in modified HBSS and mounted in agarose prior to sectioning at 300μm using a vibratome. Slices were transferred to a glass bottom culture dish coated with collagen and incubated in Slice Culture Medium (DMEM/F12 solution, supplemented with N2 solution and B27 solution without vitamin A, 5% horse serum and 5% fetal bovine serum, FGF (10ng/ml final concentration)). Brain sections were imaged using a Nikon A1plus inverted confocal microscope. For live imaging experiments, a z-stack covering 70μm was taken every 5 minutes for 17 hours. During this imaging session, brain slices were kept in a live imaging chamber maintaining 37° C with 5% CO_2_ and humidification within the chamber.

## Acknowledgements

We thank Kyle Roux and Indra Chandrasekar for help in experimental design and for sharing reagents, and Debra Silver, for reading and giving us comments on the manuscript.

## Competing interests

No competing interests declared.

## Funding

This research was funded by the National Institutes of Health, grant number P20GM103620 to LJP. Supported by a fellowship to BLM by the USD Neuroscience, Nanotechnology and Networks program through a grant from NSF (DGE-1633213).

## Supplementary material legends

**Table S1**. Table providing the sequences of crRNAs and HDR ssODNs, together with antibodies used in this study.

**Movie S1**. Overnight live imaging of endogenous EGFP-tagged LMNB1 in an E15.5 organotypic brain slice, following Breasi-CRISPR IUE at E13.5. Time in hh:mm.

## Notes

### Competing Interest Statement

The authors have declared no competing interest.

